# The Genomes of Nematode-Trapping Fungi Provide Insights into the Origin and Diversification of Fungal Carnivorism

**DOI:** 10.1101/2024.03.21.586190

**Authors:** Yani Fan, Minghao Du, Weiwei Zhang, Wei Deng, Ence Yang, Shunxian Wang, Luwen Yan, Liao Zhang, Seogchan Kang, Jacob L Steenwyk, Zhiqiang An, Xingzhong Liu, Meichun Xiang

**Affiliations:** State Key Laboratory of Mycology, Institute of Microbiology, Chinese Academy of Sciences, Beijing 100101, China; Department of Microbiology, School of Basic Medical Sciences, Peking University Health Science Center, Beijing 100191, China; University of Chinese Academy of Sciences, Beijing 100049, China; State Key Laboratory of Medicinal Chemical Biology, Key Laboratory of Molecular Microbiology and Technology of the Ministry of Education, Department of Microbiology, College of Life Science, Nankai University, Tianjin 300071, China; Department of Plant Pathology & Environmental Microbiology, Pennsylvania State University, PA 16802, USA; Howards Hughes Medical Institute and Department of Molecular and Cell Biology, University of California, Berkeley, CA 94720, USA; Texas Therapeutics Institute, The Brown Foundation Institute of Molecular Medicine, University of Texas Health Science Center, Houston, TX 77030, USA

## Abstract

Nematode-trapping fungi (NTF), most of which belong to a monophyletic lineage in Ascomycota, cannibalize nematodes and other microscopic animals, raising questions regarding the types and mechanisms of genomic changes that enabled carnivorism and adaptation to the carbon-rich and nitrogen-poor environment created by the Permian-Triassic extinction event. Here, we conducted comparative genomic analyses of 21 NTF and 21 non-NTF to address these questions. Carnivorism-associated changes include expanded genes for nematode capture, infection, and consumption (e.g., adhesive proteins, CAP superfamily, eukaryotic aspartyl proteases, and serine-type peptidases). Although the link between secondary metabolite (SM) production and carnivorism remains unclear, we found that the numbers of SM gene clusters among NTF are significantly lower than those among non-NTF. Significantly expanded cellulose degradation gene families (GH5, GH7, AA9, and CBM1) and contracted genes for carbon-nitrogen hydrolases (enzymes that degrade organic nitrogen to ammonia) are likely associated with adaptation to the carbon-rich and nitrogen-poor environment. Through horizontal gene transfer events from bacteria, NTF acquired the *Mur* gene cluster (participating in synthesizing peptidoglycan of the bacterial cell wall) and *Hyl* (a virulence factor in animals). Disruption of *MurE* reduced NTF’s ability to attract nematodes, supporting its role in carnivorism. This study provides new insights into how NTF evolved and diversified after the Permian-Triassic mass extinction event.

## Introduction

Fungi employ diverse strategies to acquire nutrients for growth and reproduction. Carnivorous nematode-trapping fungi (NTF) develop sophisticated trapping devices to capture and consume nematodes and other microscopic animals, such as amoebas, rotifers, and springtails (Pramer 1964). Although carnivorous fungi have been found in multiple phyla, more than 90% of the known NTF belong to the class Orbiliomycetes, a monophyletic lineage in Ascomycota (Yang et al. 2007). Ascomycota NTF develop adhesive traps and constricting rings to capture and consume nematodes, the most abundant soil animals (Nordbringhertz and Stalhammarcarlemalm 1978; van den Hoogen et al. 2019). The rarity of carnivorism among fungi has raised great interest in unraveling the origin and evolution of carnivorous traits.

The evolution of carnivorous Orbiliomycetes has been studied using multilocus phylogenetic analysis (Li et al. 2005; Yang et al. 2007; Yang et al. 2012). According to molecular clock estimates, fungi employing active carnivorism (forming constricting rings) and those engaging in passive carnivorism (forming adhesive traps) diverged shortly after the Permian-Triassic mass extinction event (Yang et al. 2012). This event resulted in a marked increase of dead plant material (Visscher et al. 1996), creating carbon rich and nitrogen poor environment. Barron (2003) hypothesized that NTF evolved the ability to capture nematodes to supplement nitrogen. Several NTF traits support this hypothesis. First, trap morphogenesis is induced only when free-living nematodes are present and usually requires physical contact between hyphae and nematodes (Tunlid et al. 1992; de Ulzurrun and Hsueh 2018). Second, NTF actively attract nematodes to their mycelia and hold them during trap formation (Lopez-Llorca et al. 2007; de Ulzurrun and Hsueh 2018). In addition, NTF’s high lignolytic and cellulolytic activities, which are advantageous for living in carbon-rich environments, have been well documented (Barron 1992; Barron 2003).

Comparative and evolutionary genomics has shed light on niche adaptation and the evolution of the genotype-phenotype map across the map (Watkinson 2016; Steenwyk and Rokas 2017; Murat et al. 2018; Steenwyk et al. 2019; Smith et al. 2020; Bajic and Sanchez 2020; Malar et al. 2021). For example, evolutionary dynamics of carbon and nitrogen metabolism have been illuminated from comparative genomics of Saccharomycotina yeast (Opulente et al. 2023). Similarly, the ancient origin of woody plant material degradation (i.e., lignin degradation) in mushroom forming fungi has also been charted using comparative evolutionary genomics (Floudas et al. 2012). Accordingly, comparative genome analyses between NTF and non-NTF hold promise to uncover candidate genomic changes underlying the evolution of nematode trapping capabilities.

Here, we conducted comparative and phylogenomic analyses of 21 NTF and 21 non- NTF Ascomycota species and identified candidate carnivorism-associated genomic changes, including horizontally transferred bacterial genes, and genomic adaptation to carbon-rich/nitrogen-poor environments. One of the horizontally transferred genes was disrupted, revealing an involvement in carnivorism. Transcriptome analysis of three NTF (*Drechslerella dactyloides*, *Dactylellina haptotyla* and *Arthrobotrys oligospora*) in the absence and presence of the nematode *Caenorhabditis elegans* (Fan et al. 2021; Yang et al. 2022) revealed that some candidate carnivorism-associated genes, like adhesive protein-coding genes, were up-regulated in the presence of nematodes. Together, these analyses reveal multiple dimensions of genomic changes that contributed to the evolutionary trajectory of NTF.

## Results

### Characteristics of the NTF genomes

The NTF genomes analyzed (16 *de novo* sequenced and 5 downloaded from GenBank) included 4 *Drechslerella* spp. forming mechanical constricting rings, 9 *Arthrobotrys* spp. forming 3-dimensional (3-D) adhesive nets, 8 *Dactylellina* spp. forming 2-D traps with the exception of *Da. cionopaga* (forming adhesive columns), and 7 other species forming adhesive knobs (Supplementary Table 1). Genome sizes ranged from 30.2 to 54.2 Mb (median 39.0 Mb), with the number of predicted protein-coding genes varying from 7,955 to 13,112 (Table 1) and 60.9-70.0% of the genes being annotated to encode proteins with Pfam domain(s). Although the N50 values of the *Da. entomopaga* (579 Kb), *Da. haptotyla* (177 Kb) and *Da. drechsleri* (743 Kb) genomes were much lower than those of the other NTF genomes (1.2-6.2 Mb), examination of near-universally single-copy orthologs (or BUSCO genes) indicated high gene content completeness (93.7-96.1%).

**Table 1.** Genome features of 21 nematode-trapping fungi.

| Species<sup>a</sup> | Size<br>(Mb) | N50<br>(Mb) | GC<br>(%) | No.<br>genes | Pfam<br>(%) | BUSCO<br>(%) |
| --- | --- | --- | --- | --- | --- | --- |
| *Arthrobotrys conoides* | 39.8 | 1.8 | 42.7 | 10,254 | 63.1 | 94.3 |
| *A. iridis* | 39.8 | 2.5 | 44.4 | 10,158 | 64.7 | 95.0 |
| *A. musiformis* | 40.8 | 2.0 | 44.3 | 10,510 | 63.0 | 95.1 |
| *A. oligospora* | 40.1 | 2.0 | 44.5 | 10,779 | 62.3 | 94.1 |
| *A. pseudoclavata* | 35.1 | 6.0 | 45.9 | 9,356 | 66.1 | 95.2 |
| *A. sinensis* | 40.6 | 2.3 | 45.6 | 11,240 | 62.1 | 95.5 |
| *A. sphaeroides* | 40.6 | 2.3 | 46.7 | 10,643 | 63.0 | 95.6 |
| *A. vermicola* | 40.7 | 1.6 | 45.6 | 10,501 | 63.0 | 95.7 |
| *A. flagrans* | 36.6 | 6.2 | 45 | 9927 | 68.2 | 93.4 |
| *Dactylellina cionopaga* | 47.4 | 2.1 | 43.8 | 12,524 | 61.5 | 93.8 |
| *Da. entomopaga* | 38.4 | 0.6 | 45 | 10470 | 61.0 | 94.6 |
| *Da. drechsleri* | 54.2 | 0.7 | 38.4 | 11,044 | 63.1 | 93.8 |
| *Da. haptotyla* | 38.9 | 0.2 | 45.7 | 10,353 | 65.1 | 94.5 |
| *Da. leptospora* | 36.9 | 1.2 | 44.6 | 9,986 | 65.4 | 94.5 |
| *Da. parvicollis* | 38.3 | 1.7 | 46.0 | 10,329 | 64.7 | 94.7 |
| *Da. querci* | 34.4 | 3.1 | 46.0 | 9,718 | 65.5 | 93.8 |
| *Da. tibetensis* | 36.4 | 1.9 | 45.1 | 10,165 | 65.6 | 93.7 |
| *Drechslerella brochopaga* | 35.8 | 1.9 | 50.5 | 9,044 | 67.1 | 96.1 |
| *Dr. coelobrocha* | 36.4 | 2.9 | 46.9 | 9,495 | 66.3 | 95.3 |
| *Dr. dactyloides* | 37.7 | 1.3 | 50.3 | 8,978 | 67.2 | 96.1 |
| *Dr. stenobrocha* | 30.2 | 4.8 | 50.4 | 7,955 | 70.0 | 94.5 |

<sup>a</sup>The sources of the sequenced strains and their accession numbers are shown in Supplementary Table 1.

We compared the 21 NTF genomes with the genomes of 21 non-NTF species (Ascomycota) representing diverse lifestyles, including saprophytic, mutualistic, phytopathogenic, endophytic, entomopathogenic, and nematode endoparasitic (Fig. 1A; Supplementary Tables 1 and 2). We identified 22,679 orthologous groups (OGs) among the 458,922 protein-coding genes on the 42 genomes using OrthoFinder (Supplementary Table 3). Species-specific OGs account for 0-6.3%, and 25.1-47.2% of the OGs in each species are present in all 42 genomes (Fig. 1B; Supplementary Table 4). In total, 514 OGs are NTF-specific and present in all NTF, accounting for 5.4-6.8% of the total genes in each species (Fig. 1B; Supplementary Table 5). However, 65.4% of the NTF-specific OGs could not be annotated (orphan genes). Some OGs are unique to each NTF lineage and may be associated with unique trapping strategies: 52 OGs in 3-D adhesive net-forming species, 16 OGs in 2-D adhesive net-forming species, and 31 OGs in mechanical trap-forming species.

**Fig. 1.**
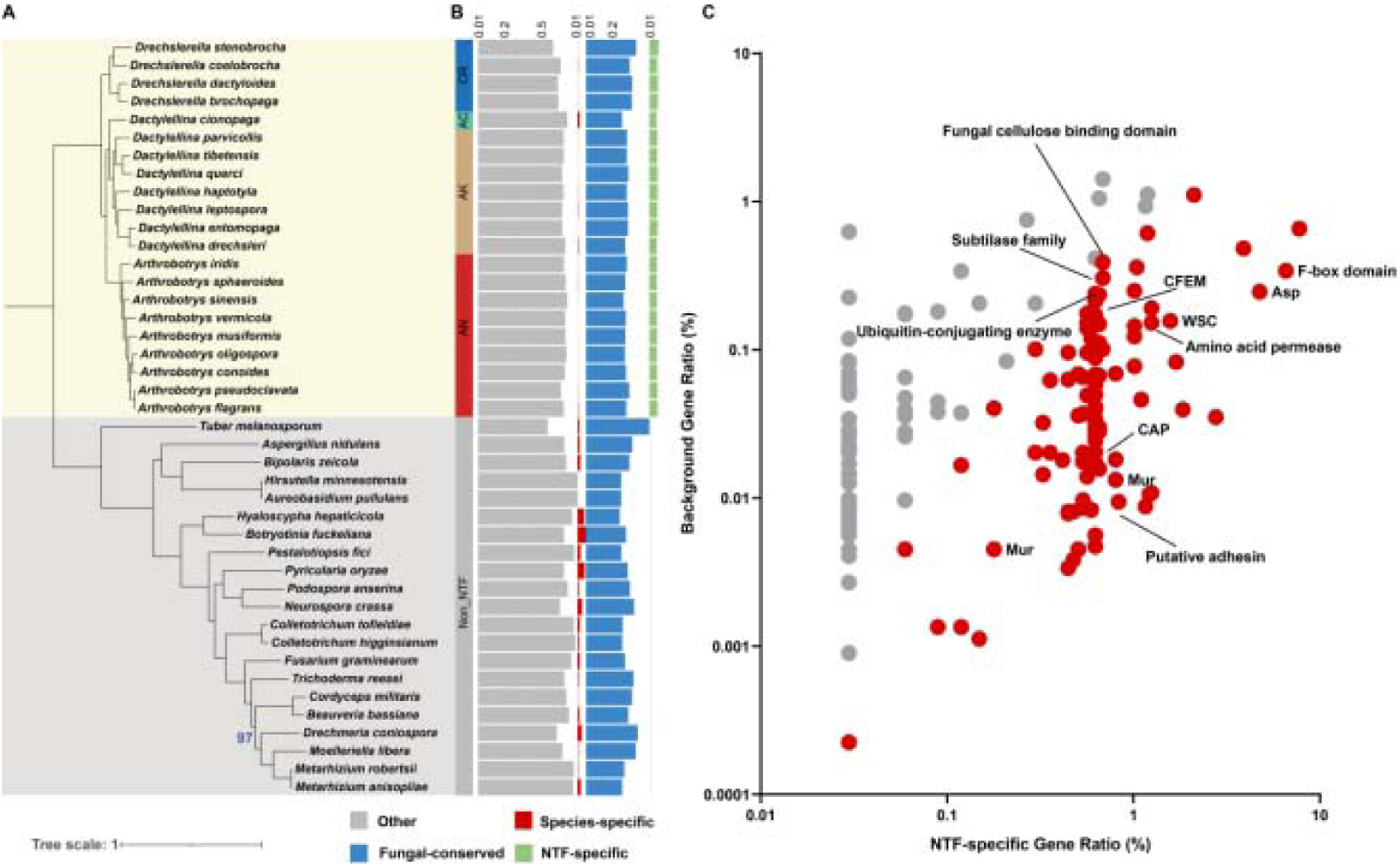
Phylogenetic relationship of the species analyzed and characteristics of the genes in individual genomes. (A) Maximum likelihood phylogeny of 21 NTF (yellow background) and 21 non-NTF (gray background) constructed using protein sequences of 704 single-copy orthologous genes present in all species. The following color scheme was used to denote different trapping devices: green (adhesive columns, AC), gold (adhesive knobs, AK), red (adhesive nets, AN), and blue (constricting rings, CR). All bootstrap values were 100 unless otherwise indicated. (B) The bar chart shows the total number of protein-coding genes in each species. The genes were classified as fungal-conserved (blue), NTF-specific (green), species-specific (red), and those present only in some fungi (gray) based on OrthoFinder group. (C) The plot shows the distribution of Pfam domains among the NTF-specific genes. Each dot represented a gene illustrated by background gene ratio (%) and NTF-specific gene ratio (%) classified by Pfam domains. Enriched Pfam domains among the NTF- specific genes (see Dataset S1, Table S3) are highlighted in red (*p*-value < 0.05, Mann-Whitney U test) and include those associated with nematode capture (CFEM domain (PF05730)), nematode infection and consumption (eukaryotic aspartyl protease (PF00026), subtilase family (PF00082) and cysteine-rich secretory protein family (PF00188)), and ubiquitination degradation of proteins such as F-box domain (PF00646), amino acid permease (PF00324) and ubiquitin-conjugating enzyme (PF00179).

Functional enrichment analysis of the NTF-specific OGs containing Pfam domain(s) showed significant enrichment of multiple gene families related to nematode capture (CFEM domain and putative adhesin) and infection and digestion [cysteine-rich secretory protein family (CAP superfamily), eukaryotic aspartyl protease (ASP), and subtilase family]. In addition, cellulose-binding domains (fungal cellulose-binding domain and WSC domains), protein ubiquitination degradation-related domains (F- box domain, ubiquitin-conjugating enzyme), amino acid permeases, and Mur proteins (involved in synthesizing peptidoglycan, a main component of the bacterial cell wall) are significantly enriched in the NTF-specific OGs (Fig. 1C; Supplementary Table 7). A search of the Non-Redundant Protein Sequence Database using the 514 NTF- specific OGs and subsequent HGTector analysis revealed 89 putative horizontal gene transfer (HGT) events. Four OGs containing different Mur protein domains and one OG with a “polysaccharide lyase family 8 domain” may participate in carnivorism. These domains exhibit significant sequence similarity to bacterial proteins but lack homologous fungal proteins (Supplementary Table 8), suggesting their horizontal gene transfer (HGT) from bacteria.

Principal component analysis (PCA) revealed significant differences in the conserved Pfam domains between NTF and non-NTF. NTF formed a discrete cluster separated from non-NTF according to PC1 (Fig. 2A). The Pfam domains that contributed the most (top 10%, 451 Pfam domains) to PC1 included 153 expanded and 298 contracted domains (Fig. 2B; Supplementary Table 9). The expanded genes include (a) those encoding extracellular proteins such as “Egh16-like virulence factor”, and “cysteine- rich secretory protein family”, (b) proteases such as “Matrixin”, protein ubiquitination degradation-related domains such as “F box”, and (c) cellulose-binding modules such as “fungal cellulose-binding domain”. In contrast, the genes for carbon-nitrogen hydrolases and secondary metabolism, such as “cytochrome P450”, “polyketide synthase dehydratase”, and “acyl transferase domain”, were contracted (Supplementary Table 9).

**Fig. 2.**
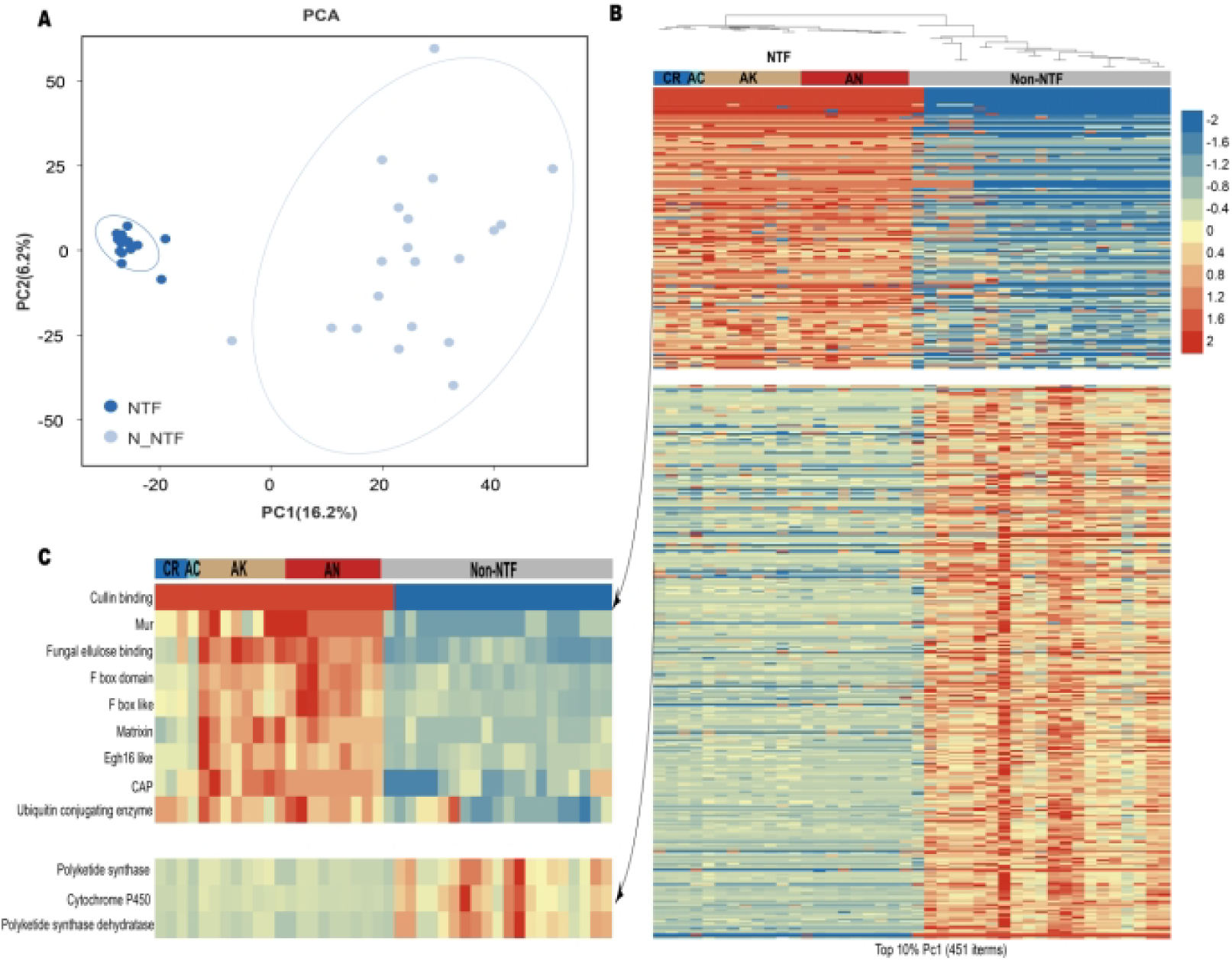
Contrasting diversification of protein domains between NTF and non-NTF. **(A)** Principal component analysis (PCA) based on the presence and number of Pfam domains in multiple orthologous groups (OGs) across NTF and non-NTF. The first two principal components account for 16.2% and 6.2% of variation, respectively. The 21 NTF are clustered together according to PC1. **(B)** Heatmaps were compiled from the Pfam data that contributed most (top 10%, 451 items). Different patterns of Pfam domain expansion and contraction are seen between NTF (yellow background) and non-NTF (gray background) based on their normalized numbers. The following color scheme was used to denote different trapping devices: green (adhesive columns, AC), gold (adhesive knobs, AK), red (adhesive nets, AN), and blue (constricting rings, CR). **(C)** Heatmaps highlighting the candidate Pfam domains related to carnivorous lifestyle selected from those shown in **B**.

### Genome changes likely associated with adaptation to carbon-rich and nitrogen-poor environments

The hypothesis that fungal carnivorism evolved in response to mass extinction was proposed (Barron 2003) but has not been tested (Yang et al. 2012). Analysis of the gene families involved in carbohydrate metabolism (Supplementary Table 10) showed that the number of carbohydrate-active enzyme (CAZyme) genes ranged from 278 to 500 in NTF (mean of 409), which is significantly lower than that in plant-associated non-NTF (endophytic: mean of 786, p = 0.0252; phytopathogenic: mean of 591, p = 0.0007; mutualistic: mean of 570, P = 0.0085) but significantly higher than that in animal parasitic non-NTF, such as entomopathogenic (mean of 352, p = 0.0027) and nematode-endoparasitic fungi (mean of 269, p = 0.0333) (Fig. 3A). Moderate expansion of the genes encoding cellulose-degrading enzymes, including GH5, GH7, and AA9, in NTF likely enhanced their cellulose-degrading capability. Moreover, the NTF genomes encode a larger set of proteins carrying one or more carbohydrate- binding modules (CBM) compared to the non-NTF genomes (Fig. 3B). CBMs, particularly CBM1 (Chundawat et al. 2021), are essential for cellulases to bind to the cellulose surface, thus enhancing the efficiency of cellulose-degrading enzymes (Espagne et al. 2008; Klosterman et al. 2011; Liu et al. 2014). The number of cellulose-degrading enzymes with CBM1 in NTF is much higher than that in non- NTF (Mann-Whitney U test) (Fig. 3C; Supplementary Table 11), a feature that likely enhanced NTF’s ability to degrade cellulose.

**Fig. 3.**
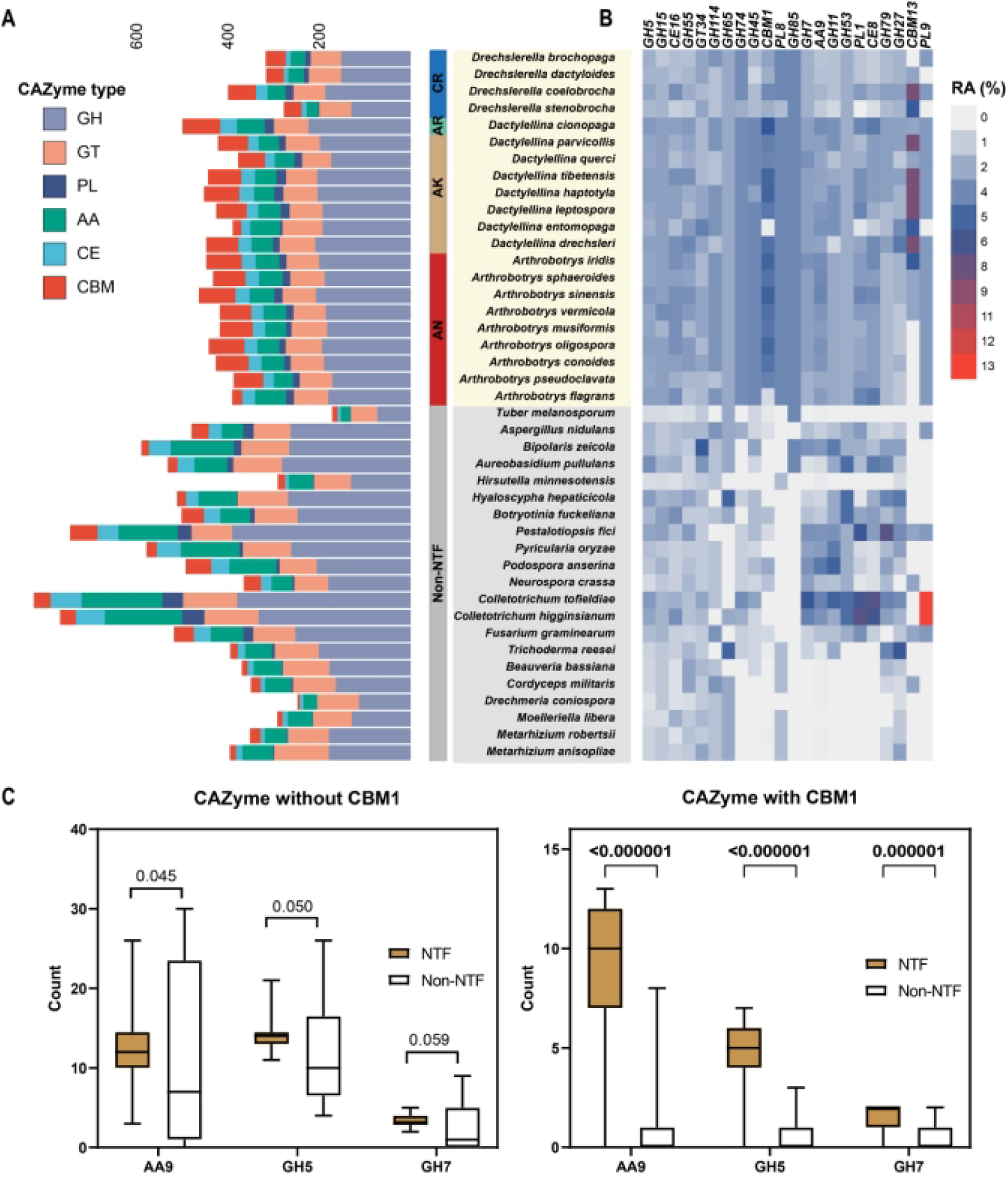
Comparative analysis of the CAZymes among NTF and non-NTF. (**A**) The bar chart shows the numbers of the following CAZymes encoded by 21 NTF (yellow background) and 21 non-NTF (gray background): glycoside hydrolases (GH), glycosyl transferases (GT), carbohydrate esterases (CE), polysaccharide lyases (PL), auxiliary activities (AA), and carbohydrate-binding modules (CBM). The following color scheme was used to denote different trapping devices: green (adhesive columns, AC), gold (adhesive knobs, AK), red (adhesive nets, AN), and blue (constricting rings, CR). (**B**) Relative abundance (RA) profiles of 13 GHs, 2 GTs, 2 CBMs, 3 PLs, 1 AAs, and 2 CEs involved in degrading cellulose. The gene number for each class was divided by the total gene number. (**C**) Comparison of the numbers of cellulose degrading enzymes with or without CBM1 between NTF (light brown) and non-NTF (white) using Mann-Whitney U test (*p*-values are shown on the graphs). The numbers of AA9, GH5, and GH7 without CBM1 (left) or with CBM1 (right) are presented.

Carnivorous fungi prey on nematodes to supplement nitrogen intake (Barron 2003; Yang et al. 2012; Lee et al. 2020). The gene family encoding carbon-nitrogen hydrolases (EC 3.5.1.-) contracted in NTF (Fig. S1; Supplementary Table 12). These hydrolases conserved among NTF (Fig. S1) belong to the nitrilase superfamily, which can break carbon-nitrogen bonds to degrade organic nitrogen compounds and produce ammonia (Pace and Brenner 2001). In contrast, many more carbon-nitrogen hydrolase coding genes were identified in entomopathogens and nematode endoparasites, suggesting that they mainly utilize protein-derived nutrients for energy (Supplementary Table 12). The genes encoding amino acid permeases are enriched in the NTF-specific OGs and contribute to amino acid transport (Supplementary Tables 7). These patterns suggest that NTF evolved to utilize organic nitrogen more efficiently by contracting carbon-nitrogen hydrolase genes (to reduce the loss of nitrogen in the form of ammonia) and gaining specific amino acid permease genes (to assimilate nitrogen in nitrogen-poor environments), supporting the hypothesis that these genomic changes were selected to help NTF adapt to nitrogen-poor conditions.

### Genome evolution putatively linked to carnivorism

NTF are expected to secrete numerous proteins that participate in capturing and consuming nematodes. We compared the predicted secreted proteins between NTF and non-NTF. The number of secreted proteins ranged from 284 to 1,654 (Fig. S2; Supplementary Tables 12 and 13). After normalization using the total number of genes for each genome, the ratio of secreted proteins encoded by NTF was significantly higher than that of non-NTF (p-value < 0.0001, Mann-Whitney U test), suggesting ancestral burst of the secreted protein repertoire.

The extracellular adhesive layer of traps is essential for nematode capture. Three types of adhesive proteins, including those containing GLEYA (PF10528), Egh16-like (PF11327), and CFEM (PF05730), were predicted to be involved in capturing nematodes (Liang et al. 2015; Ji et al. 2020; Zhang et al. 2020). These adhesive proteins accounted for more than 2% of the secreted proteins in all NTF, which is higher than that in non-NTF (Fig. 4A; Supplementary Table 14). In addition, the ratios of GLEYA domains (p-value =< 0.0001, Mann-Whitney U test) and Egh16-like domains (p-value < 0.0001, Mann-Whitney U test) in NTF are significantly higher than those in non-NTF. Gene expression patterns in three representative NTF (*Ar. oligospora*, *Da. haptotyla* and *Dr. dactyloides*) during nematode capture showed that the genes for 9.5% (2/21) of the adhesive proteins with a GLEYA domain and 15.2% (5/33) of the adhesive proteins with an Egh16-like domain were up-regulated (Supplementary Tables 15 and 17), supporting the hypothesis that expansion and up- regulation of adhesive protein-coding genes represent a genomic adaptation for a predatory lifestyle.

**Fig. 4.**
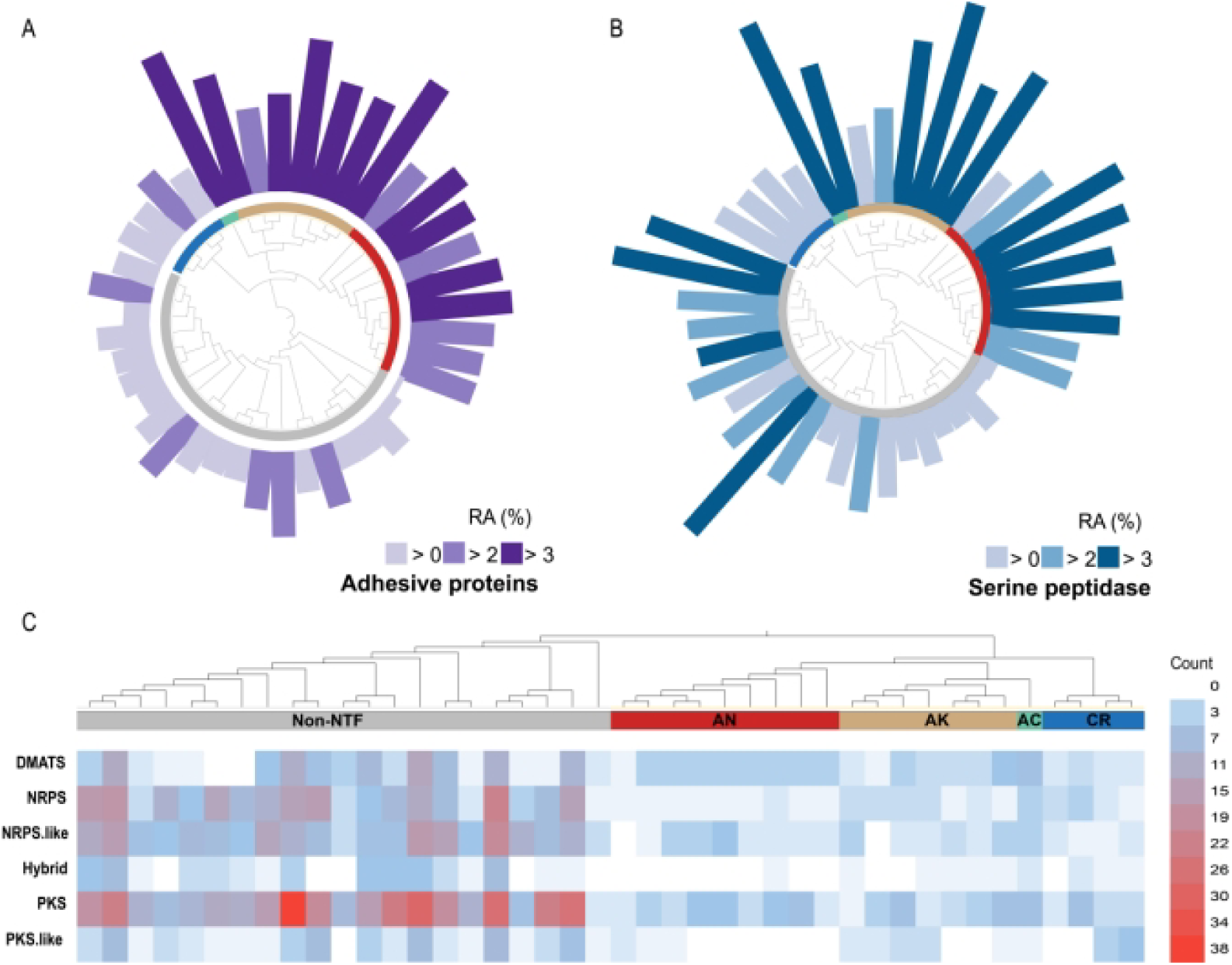
Distribution patterns of adhesive proteins, serine peptidases, and secondary metabolism clusters among NTF and non-NTF. The following color schemes were used to denote different trapping devices: blue (constricting rings, CR), red (adhesive nets, AN), green (adhesive columns, AC), and gold (adhesive knobs, AK). (**A**, **B**) Relative abundance (RA) of adhesive proteins (**A**) and serine peptidases (**B**) among the total secreted proteins encoded by each species. The numbers of adhesive proteins and serine peptidases were divided by the total number of secreted proteins to calculate their RA. (**C**) A heatmap shows the numbers of different types of secondary metabolism gene clusters in each species.

The nematode cuticle is a three-layered structure consisting mainly of collagen and noncollagenous proteins. The CAP superfamily (PF00188), composed of cysteine-rich secretory proteins, antigen 5, and pathogenesis-related 1 proteins, expanded among NTF (Supplementary Table 9). This superfamily participates in reproduction, virulence, venom toxicity, cellular defense, and immune evasion (Gibbs et al. 2008; Darwiche et al. 2016). All three NTF analyzed showed that 36.4% (4/11) of the CAP superfamily genes were up-regulated in the presence nematodes (Supplementary Tables 15 and 18), suggesting their involvement in nematode infection. In addition, the genes for serine peptidases, which are involved in nematode consumption, also significantly expanded in NTF (Fig. 4B), and 15.6% (10/64) of the genes encoding members of the subtilisin family in three NTF were up-regulated during predation (Supplementary Tables 15 and 19).

Fungi produce diverse secondary metabolites (SMs), some of which are “chemical weapons” against other organisms (Rohlfs and Churchill, 2011; Keller 2019). Significantly reduced numbers of and less conserved SM gene clusters among NTF suggest that broad capacity for SMs production is not critical for carnivorism. However, similar to other Ascomycota fungi, individual SMs may be important for organismal ecology (Raffa and Keller, 2019; Steenwyk et al. 2020a). The number of predicted SM gene clusters ranged from 6 to 23 in NTF (mean of 13.14), which is significantly lower than those of non-NTF: endophytic and phytopathogenic fungi (mean of 49.42, p < 0.0001) and entomopathogenic and nematode-endoparasitic fungi (mean of 59.29, p < 0.0001) (Fig. 4C; Supplementary Table 20). Only *T. melanosporum*, a mutualistic ectomycorrhizal fungus, has 9 clusters. The PKS gene clusters were not conserved among NTF, suggesting that this class of SMs may not be critical for carnivorism and perform species-specific roles.

Non-vertical evolution, including HGT, has been instrumental in driving the rapid adaptive evolution of fungi and has played a role in the emergence of new pathogens (Feurtey and Stukenbrock, 2018; Steenwyk et al. 2020b). Among the 89 potential HGT events observed in NTF, the HGT of *Mur* genes, which are involved in bacterial cell wall biosynthesis (Radkov et al. 2018), is notable. Top 100 blastp results using *Ar. oligospora* proteins revealed that 4 proteins with Mur domains were highly similar to MurA (53.1-69.2% identity; 90-96% coverage), MurC (65.8-72% identity; 98–99% coverage), MurD (63.7-70.2% identity; 98–99% coverage), and MurE (39.2-49.4% identity; 92–98% coverage), respectively. Maximum-likelihood phylogenetic trees were constructed using RAxML (Fig 5A, Fig. S3). The *MurE* gene was found in all NTF, suggesting its horizontal transfer before NTF diversification. The presence of introns in the NTF *MurE* gene indicates that the gene underwent eukaryotization. The function of NTF *MurE* in carnivorism was studied by disrupting the gene in *Dr. dactyloides*. Compared with the wild type, three independently isolated mutants showed reduced ability to attract *C. elegans* (p-value = 1.1e-4, 2.5e-3, 5.2e-4, two-tailed t-test, n = 5), with the attraction indices of the mutants being only 36.0% of that of the wild type (Fig. 5B).

**Fig. 5.**
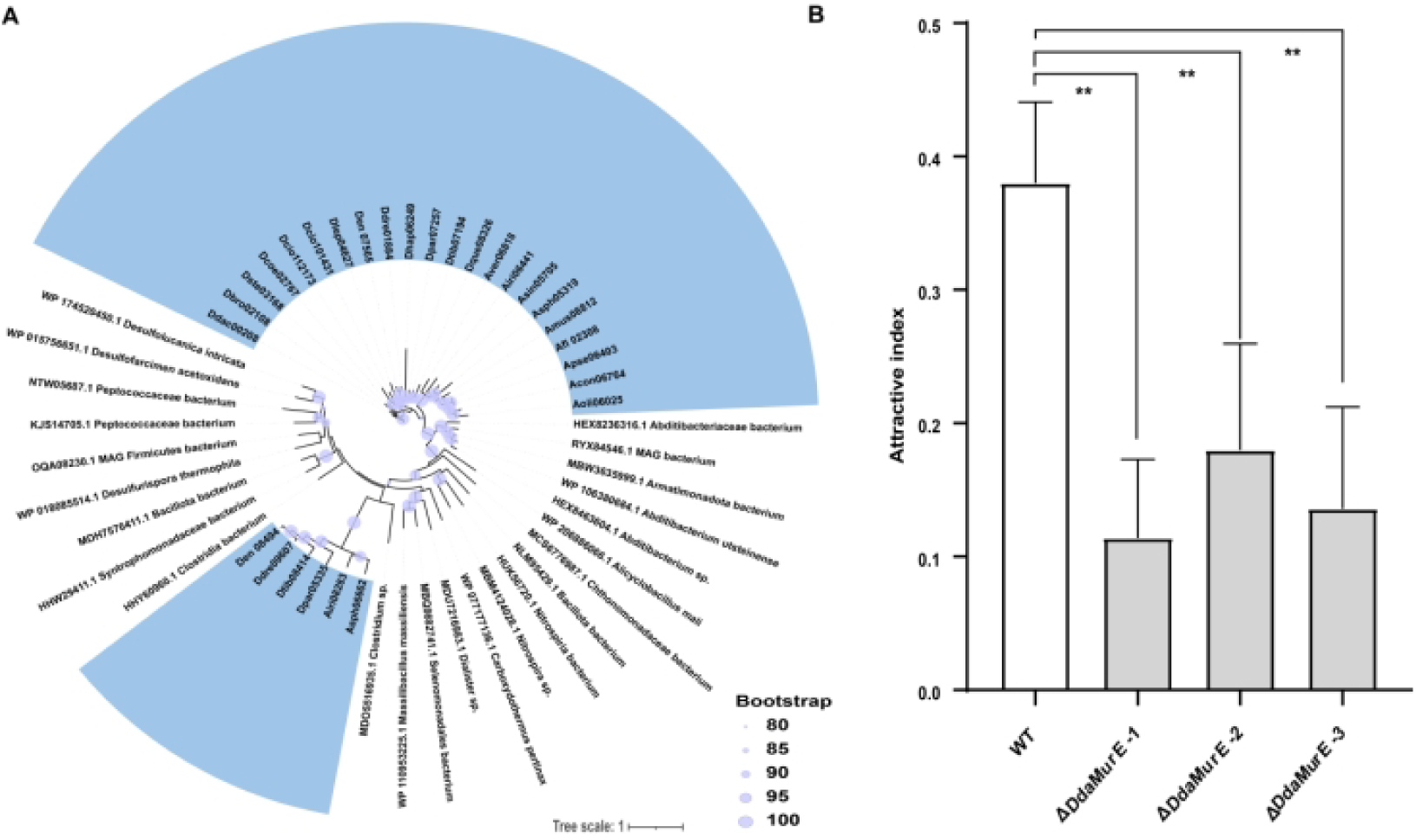
The phylogenetic relationship and potential function of *MurE*. (A) Unrooted maximum-likelihood trees based on *MurE* protein sequences were constructed using RAxML. The bacterial species (only one genome sequence selected for each species) included were chosen based on the top 100 BlastP results with the corresponding *Arthrobotrys oligospora* protein sequences as queries. All NTF species included in blue background. (B) Attractive indexes of *Drechslerella dactyloides* wild type (WT) strain and three ΔDdaMurE mutants on water agar (WA) medium. \*\**p*- value < 0.01, two-tailed t-test, n = 5.

**Fig. 6.**
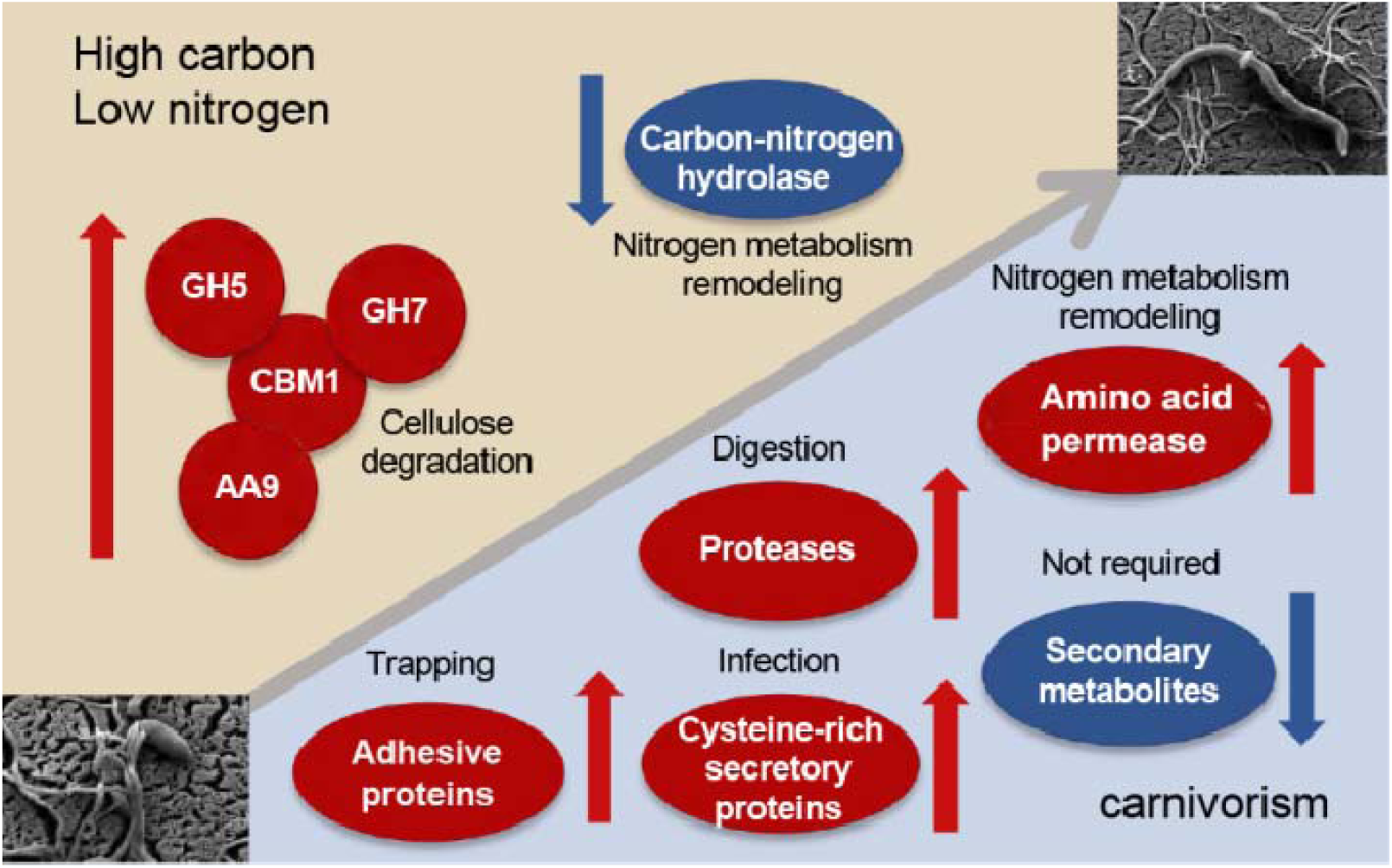
Proposed model for genomic changes associated with the evolution of NTF. To adapt to carbon-rich/nitrogen-poor environments, the genes for aminotransferase class-III and cellulose degradation enzymes expanded, whereas the genes for carbon- nitrogen hydrolase contracted. The genes encoding adhesive proteins, cysteine-rich secretory proteins, and proteases, which are likely involved in nematode capture, infection, and consumption, respectively, were expanded to support carnivorism. The number of secondary metabolite gene clusters was significantly reduced.

Analysis using CLEAN (contrastive learning-enabled enzyme annotation), which predicts enzyme function (Gregoire et al. 2023), showed that one NTF-specific OG with the polysaccharide lyase family 8 domain is closely related to Hyl, a bacterial hyaluronate lyase (EC:4.2.2.1). Bacterial Hyl is an important virulence factor employed by Gram-positive bacteria to enhance their infectivity by degrading extracellular hyaluronate and chondroitin sulfate of animal hosts (Patil et al. 2023). All NTF species encode two Hyl homologs, except for *Dr. stenobrocha* (one homolog). The maximum-likelihood phylogenetic tree of this OG showed that the two homologs belong to distinct clades (Fig. S4), suggesting that they may have originated from different bacteria through two separate HGT events. Bacterial hyaluronate lyases are typically secreted; however, their NTF homologs lack a signal peptide. Most of one Hy1 clade have putative transmembrane domains, whereas those in the other clade lack transmembrane domains (Fig. S4). Further studies are required to determine whether Hyl homologs contribute to nematode carnivorism.

## Discussion

Approximately 5 million years after the Permian-Triassic mass extinction event, the global ecosystem had stabilized and undergone extensive remodeling during the process (Ezcurra and Butler 2018; Rampino et al. 2020). The sharp increase in the biomass and diversity of taxa, including fungi, algae, and ferns, in the immediate aftermath of the plant collapse in the late Permian (Mays et al. 2020) indicated biotic recovery. The availability of new niches resulting from this mass extinction event likely accelerated innovation during recovery (Lowery and Fraass 2019).

The lifestyle of the fungi that gave rise to NTF after mass extinction remains unclear. In this study, we identified multiple genome changes—gene family gain/loss and HGT—that are associated with the evolution of NTF ecology and carnivorous lifestyle. For example, gene families that have been reported to play critical roles in trap morphogenesis (Yang et al. 2018; Yang et al. 2020; Zhang et al. 2021), such as components of the G-protein mediated signaling and ubiquitin-conjugating enzyme Ubr1 (Supplementary Table 9), have expanded. Similarly, the genes of cell-surface proteins containing the carbohydrate-binding WSC domain were up-regulated during nematode infection (Andersson et al. 2013; 2014) and were found to be significantly expanded in NTF (Supplementary Table 9). Other expanded gene families that may be linked to fungal carnivorism are those encoding adhesive proteins and proteases such as subtilase (Liang et al. 2015; Wang et al. 2015; Ji et al. 2020; Zhang et al. 2020). Overall, the patterns of gene family expansion in NTF resemble those observed in plant pathogens rather than patterns found in insect and animal pathogens (Meerupati et al. 2013).

The evolution of various trapping devices by fungi to capture nematodes as a nitrogen source was hypothesized to be an innovation crucial for NTF (Yang et al. 2012). Here we found two changes that helped optimize this mode of nitrogen acquisition. Carbon-nitrogen hydrolase genes were reduced among NTF, a change critical for efficient utilization of nitrogen resources by reducing the decomposition of organic nitrogen compounds to ammonia. Meanwhile, genes for specific amino acid permeases expanded in NTF. Some class III aminotransferase genes, which help enhance nitrogen assimilation, are up-regulated during nematode capturing (Supplementary Table 16). These patterns suggest that NTF evolved to recycle nitrogen resources efficiently by remodeling their nitrogen metabolism.

Perhaps the most strikingly, NTF have horizontally acquired multiple *Mur* genes, which are involved in bacterial cell wall peptidoglycan synthesis, from bacteria to ensnare nematodes. Disruption of the *MurE* gene indicated its involvement in attracting nematodes (Fig. 5B). Phylogenetic analysis revealed that the constituents of the *Mur* gene cluster were acquired through multiple complex HGT events (Figs. 5A and S3). Gene clusters originating from multiple HGT events have been observed in *Saccharomycotina* yeast (Gonçalves C and Gonçalves P, 2019), and our analyses suggest this may be a more widespread phenomenon in fungi. The cell wall of the nematode traps seemed different from that of hyphae (Fig. S5), suggesting that cell wall composition may play a role in trapping nematodes. How the *MurE* gene contributes to nematode attraction and whether other *Mur* genes have similar roles remain to be investigated. However, their involvement in bacterial cell wall formation raises the possibility of their involvement in modifying fungal cell wall to attract nematodes.

Our study uncovered multiple modes of genomic changes that likely influenced the evolutionary trajectory of NTF, implicating several gene families that have previously been shown to be linked to the NTF lifestyle. More importantly, our comprehensive analysis identified novel candidate genes and evolutionary processes that may underlie specific stages of predation—such as attraction and consumption. Taken together, our work established an extensive genome resource and ample hypotheses that will guide future studies to understand the origins and evolution of NTF and confirm the involvement of candidate genes associated with NTF lifestyles.

## Materials and Methods

### Strains, media, and culturing conditions

The NTF sequenced (Supplementary Table 1) were cultured on potato dextrose agar (PDA) and corn meal agar (CMA). Mycelia used for genomic DNA extraction were prepared by inoculating approximately 10^5^ spores or ten 5-mm diameter plugs from freshly grown culture on PDA into 100 mL potato dextrose broth (PDB). After shaking the cultures at 120 rpm and 28 °C for 10 days, mycelia were collected using a glass cotton filter and washed three times with distilled water.

### Whole-genome sequencing

Genomic DNA was extracted using a CTAB/SDS/Proteinase K method (Möller et al. 1992). Four genomic DNA libraries with insert sizes of 400 bp, 500 bp, 3 Kb, and 5-8 Kb, respectively were prepared. The library with 400 bp inserts was sequenced using Illumina PE250. The remaining libraries were sequenced using Illumina PE150. The resulting sequence reads were processed using Trimmomatic v0.38 to remove adapters and low-quality reads (Bolger et al. 2014). Processed reads were assembled de novo using ALLPATH-LG release 52488 (Gnerre et al. 2011). To improve completeness, draft assembled genomes were scaffolded using data from mate-pair libraries (3 Kb and 5-8 Kb) via two rounds of SSPACE-standard v3.0 (Boetzer et al. 2011). The output was used for the analyses described below. The completeness of individual genome assemblies and gene prediction was evaluated using BUSCO v4.0.2, based on Ascomycota ortholog database containing 1,706 orthologs (Boetzer et al. 2011).

### Gene prediction and functional annotation

Regions of repetitive sequence elements in the assembly were masked using RepeatMasker v4.0.7 based on a species-specific repeat library generated using RepeatModeler v4.0.7. For structural annotation, the quality of protein sequence predictions was assessed via BUSCO analysis. The BRAKER pipeline (Hoff et al. 2016) was used to test protein homology. For functional annotation, the final gene set for each species was analyzed using Pfam, PRINTS, PANTHER, SUPERFAMILY, SMART, and Gene 3D in InterProScan v5.39-77.0 (Jones et al. 2014).

The carbohydrate-active enzymes (CAZymes) were predicted by aligning all protein- coding genes in each species against the dbCAN2 database (Zhang et al. 2018) using DIAMOND v0.9.24.125, HMMER v3.0, and Hotpep. Those supported by more than two aligners were considered as CAZymes. The Secondary Metabolite Unknown Regions Finder (SMURF) was used to predict SM gene clusters (Khaldi et al. 2010). Secreted proteins were identified using multiple processes. SignalP v5.1 (Petersen et al. 2011) was initially used to predict the candidates. Subsequently, putative membrane proteins were filtered out using TMHMM v2.0 (Krogh et al. 2001). Likely mitochondrial or endoplasmic reticulum proteins were removed using WoLF PSORT, TargetP v2.0, and Deeploc v2.0 (de Castro et al. 2006; Horton et al. 2007; Armenteros et al. 2019).

### Phylogenomic data matrix construction and analysis

Concatenation is a popular method for inferring organismal histories (Steenwyk et al. 2023) and has been successfully used to infer evolutionary relationships in fungi (Li et al. 2021). Orthologous relationships among the genes in 21 NTF and 21 non-NTF were determined using OrthoFinder v2.2.6, with default settings (Emms and Kelly 2019). Single-copy genes present in all species were used for phylogenetic analysis. For each group of single-copy orthologous genes, their predicted protein sequences were aligned using MAFFT v7.453 (Katoh and Standley 2013). Aligned sequences were trimmed using the “gappyout” model in trimAl v1.2 (Capella-Gutiérrez et al. 2009), which has been demonstrated to be an efficacious approach for trimming (Tan et al. 2015; Steenwyk et al. 2020c), and concatenated. A maximum likelihood tree was generated using RAxML v8.2.12 (Stamatakis 2014) under the Q.insect+F+I+R10 model, which was automatically selected, with a discrete gamma distribution of rates across sites. 1,000 bootstrap resamplings were performed to evaluate bipartition support.

### Analysis of Pfam domains

To characterize the patterns of functional diversification in functional domains across the NTF genomes, the copy numbers of each Pfam domain in 21 NTF and 21 non- NTF were subjected to principal component analysis (PCA) using the "prcomp" function in the stat package of R v3.5.1 with default settings (R Core Team 2017). We ranked the contribution to the first principal component (PC1) in descending order and selected the top 10% to display via a heatmap using the Pheatmap v1.0.12 package of R. The number of Pfam domains for each species was normalized using zero-mean normalization, and the corresponding profiles were shown in the heatmap.

### Statistical analysis

Enrichment analysis of Pfam domains in NTF-specific genes was tested by Mann- Whitney U test (McKnight and Najab 2010). The p-values were corrected using the Bonferroni method. Comparison of the number of CAZymes involved in cellulose degradation with or without carbohydrate-binding module (CBM) between NTF and non-NTF was carried out using Mann-Whitney U test utilizing "wilcox.test" function in the stat package of R. The results were visualized using the ggplot2 package v3.3.2 of R (Wickham 2009).

### Transcriptome analysis

Published transcriptome data of *D. dactyloides* (Accession: PRJNA723922), which forms constricting rings, and *A. oligospora* (Accession: PRJNA791406), which forms adhesive nets, (Fan et al. 2021; Yang et al. 2022) were downloaded from GenBank, and transcriptome data of *Da. haptotyla*, which forms adhesive knobs, was generated by us (unpublished). Normalized read counts were used to estimate gene expression levels, and those expressed at low levels (read counts < 5) were removed. Expression levels of the genes predicted to be involved in the niche adaptation based on our comparative genome analysis were compared in the presence (24 hr) and absence (0 hr) of preys (*Caenorhabditis elegans*). Genes with padj < 0.05 and |log2 Foldchange| >1 were considered differentially expressed.

### *MurE* disruption and nematode attraction assay

The *DdaMurE* gene in *D. dactyloides* strain 29 (CGMCC3.20198) was disrupted using a published protocol (Fan et al. 2021). To determine whether the mutation affects nematode attraction, 6-mm diameter plugs (collected 0.5 cm away from the edge of plates) from 14-day-old PDA cultures of *D. dactyloides* strain 29 (WT) and three Δ*DdaMurE* mutants were placed on water agar medium, and 6-mm diameter plugs of PDA medium were used as controls. 200 nematodes were added to the central location (a circle with the radius of 0.5 cm), with 5 replicates for each sample. We employed the attractive index system described in a study by Le Saux and Queneherve (2002). The system records the degree of attraction or repulsion using numbers ranging from +2 (attraction) to -2 (repulsion). Attractive indexes were calculated 6 h after placing the plates under shade. The resulting numbers were analyzed using SPSS v20.0, and significant differences were determined by a p-value < 0.01 using Student’s t-test.

## Supporting information

Supplementary informations

## Acknowledgments

We want to thank Prof. Antonis Rokas, Department of Biological Sciences at Vanderbilt University and Prof. Yafei Mao, College of Life Science, Shanghai Jiaotong University for providing valuable suggestions. This work was supported by the National Natural Science Foundation of China (grant numbers 32020103001 and 31770065) and the Startup Fund from the Nankai University to X. L. S. K. was supported by the USDA-NIFA and Federal Appropriations (Projects PEN4655 and PEN4839). J. L. S. is a Howard Hughes Medical Institute Awardee of the Life Sciences Research Foundation.

## Author contributions

X. L., M. X. and Y. F. designed the research; Y. F., M. D., W. D., L.Y. and W. Z. performed the research; Y. F., E. Y., S. W., L. Z., M. X. and X. L. contributed new reagents/analytic tools; M. D., Y. F., W. Z. W. D. and E. Y. analyzed the data; E. Y., S. K., Z. A. and X. L supervised the data analysis; Y. F., E. Y., S. K., J. S., Z. A., M. D. and X. L. wrote the paper. All authors reviewed and approved the manuscript.

## Data availability

Data from the whole genome shotgun sequencing of 16 NTF reported in this paper are available at GenBank under the BioProject number PRJNA791178 (https://www.ncbi.nlm.nih.gov/bioproject/791178). Corresponding accession numbers are JAJTUI000000000 (*Arthrobotrys conoides*), JAJTTS000000000 (*A. iridis*), JAJTTT000000000 (*A. musiformis*), JAJTTU000000000 (*A. pseudoclavata*), JAJTTV000000000 (*A. sinensis*), JAJTTW000000000 (*A. sphaeroides*), JAJTTX000000000 (*A. vermicola*), JAJTTY000000000 (*Dactylellina cionopaga*), JAJTUB000000000 (*D. drechsleri*), JAJTUC000000000 (*D. leptospora*), JAJTUD000000000 (*D. parvicollis*), JAJTUE000000000 (*D. querci*), JAKDFA000000000 (*D. tibetensis*), JAJTUF000000000 (*Drechslerella brochopaga*), JAJTUH000000000 (*Dr. coelobrocha*), and JAJTUG000000000 (*Dr. dactyloides*).

## References

Andersson KM, Kumar D, Bentzer J, Friman E, Ahren D, Tunlid A. 2014. Interspecific and host-related gene expression patterns in nematode-trapping fungi. BMC Genomics 15:968.

Andersson KM, Meerupati T, Levander F, Friman E, Ahren D, Tunlid A. 2013. Proteome of the nematode-trapping cells of the fungus *Monacrosporium haptotylum*. Appl Environ Microb. 79:4993–5004.

Armenteros JJA, Salvatore M, Emanuelsson O, Winther O, von Heijne G, Elofsson A, Nielsen H. 2019. Detecting sequence signals in targeting peptides using deep learning. Life Sci Alliance 2:e201900429.

Bajic D, Sanchez A. 2020. The ecology and evolution of microbial metabolic strategies. Curr Opin Biotechnol. 62:123–128.

Barron GL. 1992. Lignolytic and cellulolytic fungi as predators and parasites. In: Carroll GC, Wicklow DT. (eds). The fungal Community: Its Organization and Role in the Ecosystem. Abingdon: Routledge. p. 311–326.

Barron GL. 2003. Predatory fungi, wood decay, and the carbon cycle. Biodiversity 4:3–9.

Barreteau H, Kovac A, Boniface A, Sova M, Gobec S, Blanot D. 2008. Cytoplasmic steps of peptidoglycan biosynthesis. FEMS Microbiol Rev 32(2):168–207.

Boetzer M, Henkel CV, Jansen HJ, Butler D, Pirovano W. 2011. Scaffolding pre-assembled contigs using SSPACE. Bioinformatics 27:578–579.

Bolger AM, Lohse M, Usadel B. 2014. Trimmomatic: a flexible trimmer for Illumina sequence data. Bioinformatics 30:2114–2120.

Bugg TD, Braddick D, Dowson CG, Roper DI. 2011. Bacterial cell wall assembly: still an attractive antibacterial target. Trends Biotechnol 29(4):167–73.

Capella-Gutiérrez S, Silla-Martínez JM, Gabaldón T. 2009. trimAl: a tool for automated alignment trimming in large-scale phylogenetic analyses. Bioinformatics 25:1972–1973.

Chundawat S, Nemmaru B, Hackl M, Brady S, Hilton M, Johnson M, Chang S, Lang M, Hyun H, Lee S-H, et al. 2021. Molecular origins of reduced activity and binding commitment of processive cellulases and associated carbohydrate- binding proteins to cellulose III. J Biol Chem. 296:100431.

Darwiche R, Kelleher A, Hudspeth EM, Schneiter R, Asojo OA. 2016. Structural and functional characterization of the CAP domain of pathogen-related yeast 1 (Pry1) protein. Sci Rep. 6:28838.

de Castro E, Sigrist CJ, Gattiker A, Bulliard V, Langendijk-Genevaux PS, Gasteiger E, Bairoch A, Hulo N. 2006. ScanProsite: detection of PROSITE signature matches and ProRule-associated functional and structural residues in proteins. Nucleic Acids Res. 34:W362–W365.

de Ulzurrun GVD, Hsueh YP. 2018. Predator-prey interactions of nematode-trapping fungi and nematodes: both sides of the coin. Appl Microbiol Biotechnol. 102:3939–3949.

de Vries RP, Makela MR. 2020. Genomic and postgenomic diversity of fungal plant biomass degradation approaches. Trends Microbiol. 28:487–499.

Emms DM, Kelly S. 2019. OrthoFinder: phylogenetic orthology inference for comparative genomics. Genome Biol. 20:238.

Espagne E, Lespinet O, Malagnac F, Da Silva C, Jaillon O, Porcel BM, Couloux A, Aury JM, Segurens B, Poulain J, et al. 2008. The genome sequence of the model ascomycete fungus *Podospora anserina*. Genome Biol. 9:R77.

Ezcurra MD, Butler RJ. 2018. The rise of the ruling reptiles and ecosystem recovery from the Permo-Triassic mass extinction. Proc R Soc B. 285:20180361.

Fan YN, Zhang WW, Chen Y, Xiang MC, Liu XZ. 2021. DdaSTE12 is involved in trap formation, ring inflation, conidiation, and vegetative growth in the nematode-trapping fungus *Drechslerella dactyloides*. Appl Microbiol Biotechnol. 105:7379–7393.

Feurtey A, Stukenbrock Eva H. 2018. Interspecific gene exchange as a driver of adaptive evolution in fungi. Ann Rev Microbiol 72: 377–398.

Floudas D, Binder M, Riley R, Barry K, Blanchette RA, Henrissat B, Martínez AT et al. The Paleozoic origin of enzymatic lignin decomposition reconstructed from 31 fungal genomes. Science. 2012:336(6089):1715-1719.

Gibbs GM, Roelants K, O’Bryan MK. 2008. The CAP superfamily: cysteine-rich secretory proteins, antigen 5, and pathogenesis-related 1 proteins-roles in reproduction, cancer, and immune defense. Endocr Rev. 29:865–897.

Gregoire, J.M., Zhou, L., Haber, J.A. 2023. Combinatorial synthesis for AI-driven materials discovery. Nat Synth. 2:493–504.

Gnerre S, MacCallum I, Przybylski D, Ribeiro FJ, Burton JN, Walker BJ, Sharpe T, Hall G, Shea TP, Sykes S, et al. 2011. High-quality draft assemblies of mammalian genomes from massively parallel sequence data. Proc Natl Acad Sci U S A. 108:1513–1518.

Gonçalves C, Gonçalves P. 2019. Multilayered horizontal operon transfers from bacteria reconstruct a thiamine salvage pathway in yeasts. Proc Natl Acad Sci U S A. 116(44):22219–22228.

Hittinger CT, Rokas A. 2019. Extensive loss of cell-cycle and DNA repair genes in an ancient lineage of bipolar budding yeasts. PLoS Biol 17(5):e3000255.

Hoff KJ, Lange S, Lomsadze A, Borodovsky M, Stanke M. 2016. BRAKER1: unsupervised RNA-seq-based genome annotation with GeneMark-ET and AUGUSTUS. Bioinformatics 32:767–769.

Horton P, Park KJ, Obayashi T, Fujita N, Harada H, Adams-Collier CJ, Nakai K. 2007. WoLF PSORT: protein localization predictor. Nucleic Acids Res. 35:W585–W587.

Ji XL, Yu ZF, Yang JK, Xu JP, Zhang Y, Liu SQ, Zou CG, Li J, Liang LM, Zhang KQ. 2020. Expansion of adhesion genes drives pathogenic adaptation of nematode- trapping fungi. iScience 23:101057.

Jones P, Binns D, Chang HY, Fraser M, Li W, McAnulla C, McWilliam H, Maslen J, Mitchell A, Nuka G, et al. 2014. InterProScan 5: genome-scale protein function classification. Bioinformatics 30:1236–1240.

Katoh K, Standley DM. 2013. MAFFT multiple sequence alignment software version 7: improvements in performance and usability. Mol Biol Evol. 30:772–780.

Keller NP. 2019. Fungal secondary metabolism: regulation, function and drug discovery. Nat Rev Microbiol 17:167–180.

Khaldi N, Seifuddin FT, Turner G, Haft D, Nierman WC, Wolfe KH, Fedorova ND. 2010. SMURF: genomic mapping of fungal secondary metabolite clusters. Fungal Genet Biol. 47:736–741.

Klosterman SJ, Subbarao KV, Kang SC, Veronese P, Gold SE, Thomma BPHJ, Chen ZH, Henrissat B, Lee YH, Park J, et al. 2011. Comparative genomics yields insights into niche adaptation of plant vascular wilt pathogens. PLoS Pathog. 7:e1002137.

Krogh A, Larsson B, von Heijne G, Sonnhammer EL. 2001. Predicting transmembrane protein topology with a hidden Markov model: application to complete genomes. J Mol Biol. 305:567–580.

Le Saux R, Queneherve P. 2002. Differential chemotactic responses of two plant- parasitic nematodes, *Meloidogyne incognita* and *Rotylenchulus reniformis*, to some inorganic ions. Nematology 4:99–105.

Lee CH, Chang HW, Yang CT, Wali N, Shie JJ, Hsueh YP. 2020. Sensory cilia as the Achilles heel of nematodes when attacked by carnivorous mushrooms. Proc Natl Acad Sci U S A. 117:6014–6022.

Li Y, Hyde KD, Jeewon R, Cai L, Vijaykrishna D, Zhang KQ. 2005. Phylogenetics and evolution of nematode-trapping fungi (Orbiliales) estimated from nuclear and protein coding genes. Mycologia. 97:1034–1046.

Li Y, Steenwyk JL, Chang Y, Wang Y, James TY, Stajich JE, Spatafora JW, Groenewald M, Dunn CW, Hittinger CT, Shen XX, Rokas A. 2021. A genome- scale phylogeny of the kingdom Fungi. Cur Biol. 31(8):1653–1665.e5.

Liang LM, Shen RF, Mo YY, Yang JK, Ji XL, Zhang KQ. 2015. A proposed adhesin AoMad1 helps nematode-trapping fungus *Arthrobotrys oligospora* recognizing host signals for life-style switching. Fungal Genet Biol. 81:172–181.

Liu KK, Zhang WW, Lai YL, Xiang MC, Wang XN, Zhang XY, Liu XZ. 2014. *Drechslerella stenobrocha* genome illustrates the mechanism of constricting rings and the origin of nematode predation in fungi. BMC Genomics 15:114.

Lopez-Llorca LV, Maciá-Vicente JG, Jansson HB. 2007. Mode of action and interactions of nematophagous fungi. In: Ciancio A, Mukerji KG. (eds). Integrated Management and Biocontrol of Vegetable and Grain Crops Nematodes. Dordrecht: Springer. p. 43–59.

Lowery CM, Fraass AJ. 2019. Morphospace expansion paces taxonomic diversification after end Cretaceous mass extinction. Nat Ecol Evol. 3:900–904.

Marques DA, Jones FC, Di Palma F, Kingsley DM, Reimchen TE. 2018. Experimental evidence for rapid genomic adaptation to a new niche in an adaptive radiation. Nat Ecol Evol. 2:1128–1138.

Malar CM, Krüger M, Krüger C, Wang Y, Stajich JE, Keller J, Chen ECH, Yildirir G, Villeneuve-Laroche M, Roux C, Delaux PM, Corradi N. 2021. The genome of *Geosiphon pyriformis* reveals ancestral traits linked to the emergence of the arbuscular mycorrhizal symbiosis. Cur Biol. 31(7):1570–1577.e4.

Mays C, Vajda V, Frank TD, Fielding CR, Nicoll RS, Tevyaw AP, McLoughlin S. 2020. Refined Permian-Triassic floristic timeline reveals early collapse and delayed recovery of south polar terrestrial ecosystems. Geol Soc Am Bull. 132:1489–1513.

McKnight PE, Najab J. 2010. Mann-Whitney U Test. In: Weiner IB, Craighead WE, (eds). The Corsini Encyclopedia of Psychology. p. 1–1.

Meerupati T, Andersson KM, Friman E, Kumar D, Tunlid A, Ahren D. 2013. Genomic mechanisms accounting for the adaptation to parasitism in nematode-trapping fungi. PLoS Genet. 9:e1003909.

Milewski S, Gabriel I, Olchowy J. 2006. Enzymes of UDP-GlcNAc biosynthesis in yeast. Yeast. 23(1):1–14.

Möller EM, Bahnweg G, Sandermann H, Geiger HH. 1992. A simple and efficient protocol for isolation of high molecular weight DNA from filamentous fungi, fruit bodies, and infected plant tissues. Nucleic Acids Res. 20:6115–6116.

Murat C, Payen T, Noel B, Kuo A, Morin E, Chen J, Kohler A, Krizsán K, Balestrini R, Da Silva C, et al. 2018. Pezizomycetes genomes reveal the molecular basis of ectomycorrhizal truffle lifestyle. Nat Ecol Evol, 2(12):1956–1965.

Nordbring-Hertz B, Jansson HB, Tunlid A. 2006. Nematophagous fungi. In: Encyclopedia of Life Sciences. Chichester: John Wiley & Sons, Ltd. p. 1–11.

Nordbringhertz B, Stalhammarcarlemalm M. 1978. Capture of nematodes by *Arthrobotrys oligospora*, an electron microscope study. Can J Bot -Rev Can Bot. 56:1297–1307.

Opulente DA, Leavitt LaBella A, Harrison MC, et al. Genomic and ecological factors shaping specialism and generalism across an entire subphylum. Preprint. bioRxiv. 2023.06.19.545611.

Pace HC, Brenner C. 2001. The nitrilase superfamily: classification, structure and function. Genome Biol. 2:1–9.

Patil, S.P., Dalal, K.S., Shirsath, L.P., Chaudhari, B.L. 2023. Microbial hyaluronidase: its production, purification and applications. In: Verma P. (ed) Industrial Microbiology and Biotechnology. Springer, Singapore.

Petersen TN, Brunak S, von Heijne G, Nielsen H. 2011. SignalP 4.0: discriminating signal peptides from transmembrane regions. Nat Methods 8:785–786.

Pramer D. 1964. Nematode-trapping fungi. Science 144:382–388.

R Core Team. 2017. R: a language and environment for statistical computing. Vienna: R Foundation for Statistical Computing.

Radkov AD, Hsu YP, Booher G, VanNieuwenhze MS. 2018. Imaging bacterial cell wall biosynthesis. Annu Rev Biochem. 87:991–1014.

Raffa N, Keller NP. 2019. A call to arms: Mustering secondary metabolites for success and survival of an opportunistic pathogen. PLoS Pathog 15(4), e1007606.

Rampino MR, Eshet-Alkalai Y, Koutavas A, Rodriguez S. 2020. End-Permian stratigraphic timeline applied to the timing of marine and non-marine extinctions. Palaeoworld 29:577–589.

Rankin CH. 2006. Nematode behavior: The taste of success, the smell of danger! Cur Biol. 16(3): R89–R91.

Rohlfs M, Churchill ACL. Fungal secondary metabolites as modulators of interactions with insects and other arthropods. Fungal Genet Biol 2011, 48(1):23–34.

Smith SD, Pennell MW, Dunn CW, Edwards SV. Phylogenetics is the New Genetics (for Most of Biodiversity). Trends Ecol Evol. 2020:35(5):415–425.

Stamatakis A. 2014. RAxML version 8: a tool for phylogenetic analysis and post- analysis of large phylogenies. Bioinformatics 30:1312–1313.

Steenwyk JL, Rokas A. 2017. Extensive copy number variation in fermentation- related genes among *Saccharomyces cerevisiae* wine strains. G3 (Bethesda) 7(5):1475-1485.

Steenwyk JL, Mead ME, Knowles SL, Raja HA, Roberts CD, Bader O, Houbraken J, Goldman GH, Oberlies NH, Rokas A. 2020a. Variation Among Biosynthetic Gene Clusters, Secondary Metabolite Profiles, and Cards of Virulence Across Aspergillus Species. Genetics 216(2), 481–497.

Steenwyk JL, Opulente DA, Kominek J, Shen XX, Zhou X, Labella AL, Bradley NP, Eichman BF, Čadež N, Libkind D, DeVirgilio J, Hulfachor AB, Kurtzman CP, Steenwyk JL, Lind AL, Ries LNA, Dos Reis TF, Silva LP, Almeida F, Bastos RW, Fraga da Silva TFC, Bonato VLD, Pessoni AM, Rodrigues F, Raja HA, Knowles SL, Oberlies NH, Lagrou K, Goldman GH, Rokas A. 2020b. Pathogenic allodiploid hybrids of *Aspergillus* fungi. Cur Biol. 30(13):2495–2507.e7.

Steenwyk JL, Buida TJ III, Li Y, Shen X-X, Rokas A. 2020c. ClipKIT: A multiple sequence alignment trimming software for accurate phylogenomic inference. PLoS Biol 18(12): e3001007.

Steenwyk JL, Li Y, Zhou X, Shen XX, Rokas A. 2023. Incongruence in the phylogenomics era. Nat Rev Genet. 24(12):834–850.

Tan G, Muffato M, Ledergerber C, Herrero J, Goldman N, Gil M, Dessimoz C. 2015. Current Methods for Automated Filtering of Multiple Sequence Alignments Frequently Worsen Single-Gene Phylogenetic Inference. Syst Biol 64(5):778–791.

Thomson N, Bentley S, Holden M, Parkhill J. 2003. Genome watch-fitting the niche by genomic adaptation. Nat Rev Microbiol. 1:92–93.

Tunlid A, Jansson HB, Nordbringhertz B. 1992. Fungal attachment to nematodes. Mycol Res. 96:401–412.

van den Hoogen J, Geisen S, Routh D, Ferris H, Traunspurger W, Wardle DA, de Goede RGM, Adams BJ, Ahmad W, Andriuzzi WS, et al. 2019. Soil nematode abundance and functional group composition at a global scale. Nature 572:194–198.

Visscher H, Brinkhuis H, Dilcher DL, Elsik WC, Eshet Y, Looy CV, Rampino MR, Traverse A. 1996. The terminal Paleozoic fungal event: evidence of terrestrial ecosystem destabilization and collapse. Proc Natl Acad Sci U S A. 93:2155–2158.

Wang R, Wang J, Yang XY. 2015. The extracellular bioactive substances of *Arthrobotrys oligospora* during the nematode-trapping process. Biol Control. 86: 60–65.

Wang SX & Liu XZ. 2023. Tools and basic procedures of gene manipulation in nematode-trapping fungi. Mycology, 14(2): 75–90.

Wang X, Li GH, Zou CG, Ji XL, Liu T, Zhao PJ, Liang LM, Xu JP, An ZQ, Zheng X, et al. 2014. Bacteria can mobilize nematode-trapping fungi to kill nematodes. Nat. Commun. 5:5776.

Watkinson SC. 2016. Molecular ecology. In: Watkinson SC, Boddy L, Money NP, (eds). The Fungi 3rd edition. Boston: Academic Press. p. 189–203.

Wickham H. 2009. ggplot2: Elegant Graphics for Data Analysis. New York: Springer.

Yang CT, Vidal-Diez de Ulzurrun G, Gonçalves A, Lin HC, Chang CW, Huang TY, Chen SA, Lai CK, Tsai I, Schroeder F, et al. 2020. Natural diversity in the predatory behavior facilitates the establishment of a robust model strain for nematode-trapping fungi. Proc Natl Acad Sci U S A. 117:201919726.

Yang EC, Xu LL, Yang Y, Zhang XY, Xiang MC, Wang CS, An ZQ, Liu XZ. 2012. Origin and evolution of carnivorism in the Ascomycota (fungi). Proc Natl Acad Sci U S A. 109:10960–10965.

Yang L, Li XM, Bai N, Yang XW, Zhang KQ, Yang JK. 2022. Transcriptomic analysis reveals that Rho GTPases regulate trap development and lifestyle transition of the nematode-trapping fungus *Arthrobotrys oligospora*. Microbiol Spectr. 10:e0175921.

Yang XW, Ma N, Yang L, Zheng YQ, Zhen ZY, Li Q, Xie MH, Li J, Zhang KQ, Yang JK. 2018. Two Rab GTPases play different roles in conidiation, trap formation, stress resistance, and virulence in the nematode-trapping fungus *Arthrobotrys oligospora*. Appl Microbiol Biotechnol. 102:4601–4613.

Yang Y, Yang EC, An ZQ, Liu XZ. 2007. Evolution of nematode-trapping cells of predatory fungi of the Orbiliaceae based on evidence from rRNA-encoding DNA and multiprotein sequences. Proc Natl Acad Sci U S A. 104:8379–8384.

Zhang H, Yohe T, Huang L, Entwistle S, Wu P, Yang Z, Busk PK, Xu Y, Yin Y. 2018. dbCAN2: a meta server for automated carbohydrate-active enzyme annotation. Nucleic Acids Res. 46:W95–W101.

Zhang W, Liu DD, Yu ZC, Hou B, Fan Y, Li ZH, Shang SJ, Qiao YD, Fu JT, Niu JK, et al. 2020. Comparative genome and transcriptome analysis of the nematode- trapping fungus *Duddingtonia flagrans* reveals high pathogenicity during nematode infection. Biol Control. 143:104159.

Zhang WW, Chen J, Fan YN, Hussain M, Liu XZ, Xiang MC. 2021. The E3-ligase AoUBR1 in N-end rule pathway is involved in the vegetative growth, secretome, and trap formation in *Arthrobotrys oligospora*. Fungal Biol. 125(7).

