## Supplementary informations for "The Genomes of Nematode-Trapping Fungi Provide Insights into the Origin and Diversification of Fungal Carnivorism"

**This PDF file includes:**

Figures S1 to S5

Supplementary Tables 1 to 20

**The following supplementary material is presented in a separate file:**

Dataset S1 (XLSX file)

**Supplementary Information**

**Supplementary Figures**

**Fig. S1**: Maximum likelihood tree based on the carbon-nitrogen hydrolases encoded by 42 species.

**Fig. S2**: The number of secreted protein genes in the 21 NTF and 21 non-NTF genomes.

**Fig. S3**: Phylogenetic trees of the three Mur proteins encoded by NTF and selected bacteria.

**Fig. S4**: Maximum likelihood tree based on the Hyl proteins encoded by 21 NTF and selected bacteria.

**Fig. S5:** Cell wall of NTF hyphae and traps stained with Concanavalin A (ConA).

**Supplementary Tables (shown in the Dataset S1 (XLSX file)**

**Supplementary Table 1** shows detailed information about the nematode-trapping fungi (NTF) analyzed in this study.

**Supplementary Table 2** shows the characteristics of the non-NTF genomes analyzed in this study.

**Supplementary Table 3** shows 433,288 fungal genes with Pfam annotation in corresponding OGs and the geneID consisting of sampleID and five-digit numbers.

**Supplementary Table 4** shows the corresponding OG type for each OGID, and the four types of OGs, including fungal-conserved, NTF-specific, species-specific, and other OGs.

**Supplementary Table 5** shows the NTF-specific OGs conserved across the 21 NTF analyzed and the total number of genes in each species. NTF forming different trapping devices are labelled: light purple (AN), yellow (AC), light green (AK), and light red (CR). Light gray denotes non-NTF.

**Supplementary Table 6** shows specific OGs shared by NTF species forming specific trap devices and the number of OGs. The NTF forming different trapping devices are labelled as noted in Supplementary Table 5.

**Supplementary Table 7** shows the enriched Pfam domains among the NTF-specific genes.

**Supplementary Table 8** shows all predicted horizontally transferred genes among the NTF-specific genes, including those putatively involved in peptidoglycan synthesis.

**Supplementary Table 9** shows the number of Pfam domains that contributed the most (top 10%) to PC1 in NTF and non-NTF. NTF forming different trapping devices are labelled: light purple (adhesive nets, AN), yellow (adhesive columns, AC), light green (adhesive knobs, AK), and light red (constricting rings, CR). Light gray denotes non-NTF.

**Supplementary Table 10** shows the number of CAZymes in NTF and non-NTF, including glycoside hydrolases (GHs), glycosyl transferases (GTs), polysaccharide lyases (PLs), carbohydrate esterases (CEs), auxiliary activities (AAs), and CBMs.

**Supplementary Table 11** demonstrates the number of CAZymes appended with or without CBM, separately, in NTF and non-NTF, including GH5, GH7, GH10, GH11, GH13, GH16, GH18, GH71, GH72, GH74, GH78, CE1, CE15, CE16, and AA9.

**Supplementary Table 12** shows the number of predicted serine peptidases, secreted proteins, carbon-nitrogen hydrolases, and class III aminotransferases encoded by NTF and non-NTF.

**Supplementary Table 13** shows the predicted secreted proteins with Pfam annotations encoded by NTF and non-NTF.

**Supplementary Table 14** shows the numbers of three types of secreted adhesive proteins encoded by NTF and non-NTF, including GLEYA domain appended proteins, CFEM domain appended proteins, and Egh16-like virulence factors.

**Supplementary Tables S15-S19** contain transcriptome data generated from *Da. haptotyla* forming AK, *A. oligospora* forming AN, and *Dr. stenobrocha* forming CR.

**Supplementary Table 15** summarizes the number of carnivorism-related genes un-regulated in the presence of *C.* elegans. Their functions are related to nematodes adhesion, nematodes infection, nematodes digesting, and nitrogen utilization.

**Supplementary Table 16** shows the expression level of nitrogen utilization-related genes (those encoding carbon-nitrogen hydrolase and aminotransferase) in three NTF species.

**Supplementary Table 17** shows the expression level of predicted adhesive protein-encoding genes in three NTF species.

**Supplementary Table 18** shows the expression level of cysteine-rich secretory protein family members in three NTF species.

**Supplementary Table 19** shows the expression level of nematode digestion-related genes (those encoding eukaryotic aspartyl protease and subtilase) in three NTF species.

**Supplementary Table 20** shows the number of secondary metabolite (SM) gene clusters in NTF and non-NTF. Six types of SM gene clusters, including dimethylallyltryptophan synthase (DMATS), non-ribosomal peptides synthetase (NRPS), NRPS-like, polyketides synthetase (PKS), PKS-like, and NRPS-PKS-hybrid, are shown.

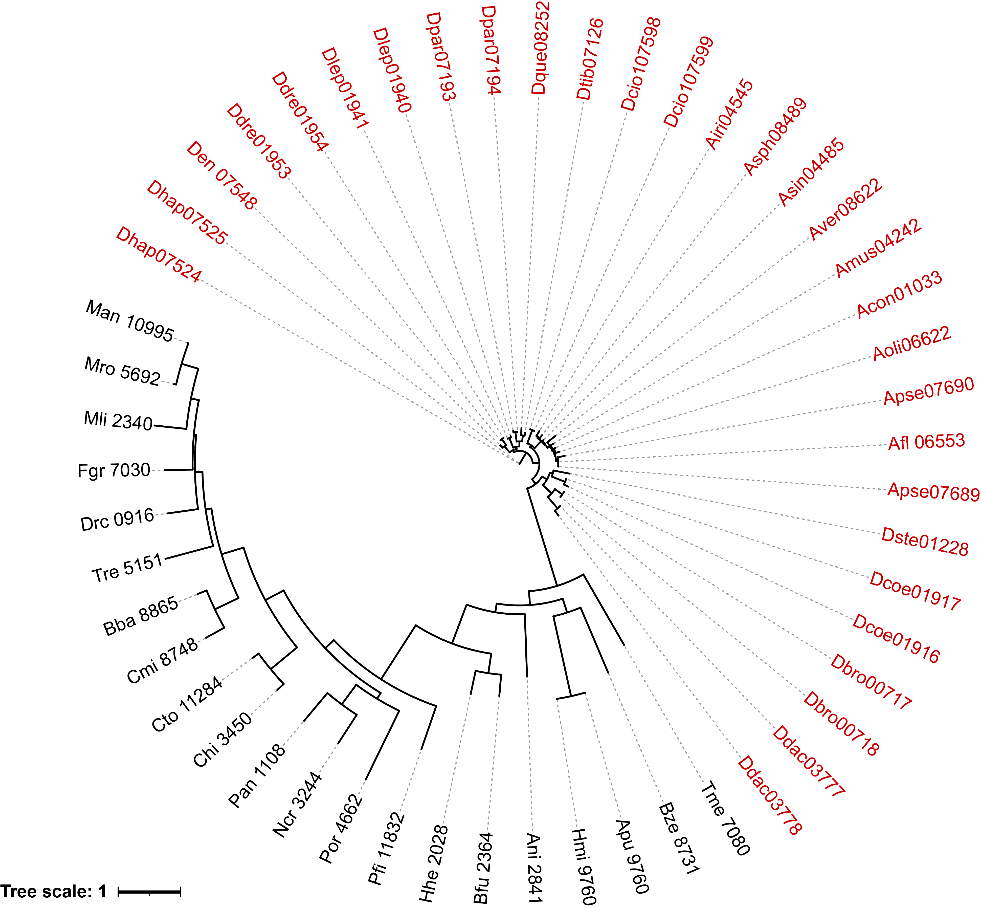

**Fig. S1: Maximum likelihood tree based on the carbon-nitrogen hydrolases encoded by 42 species.** The proteins encoded by NTF are marked in red.

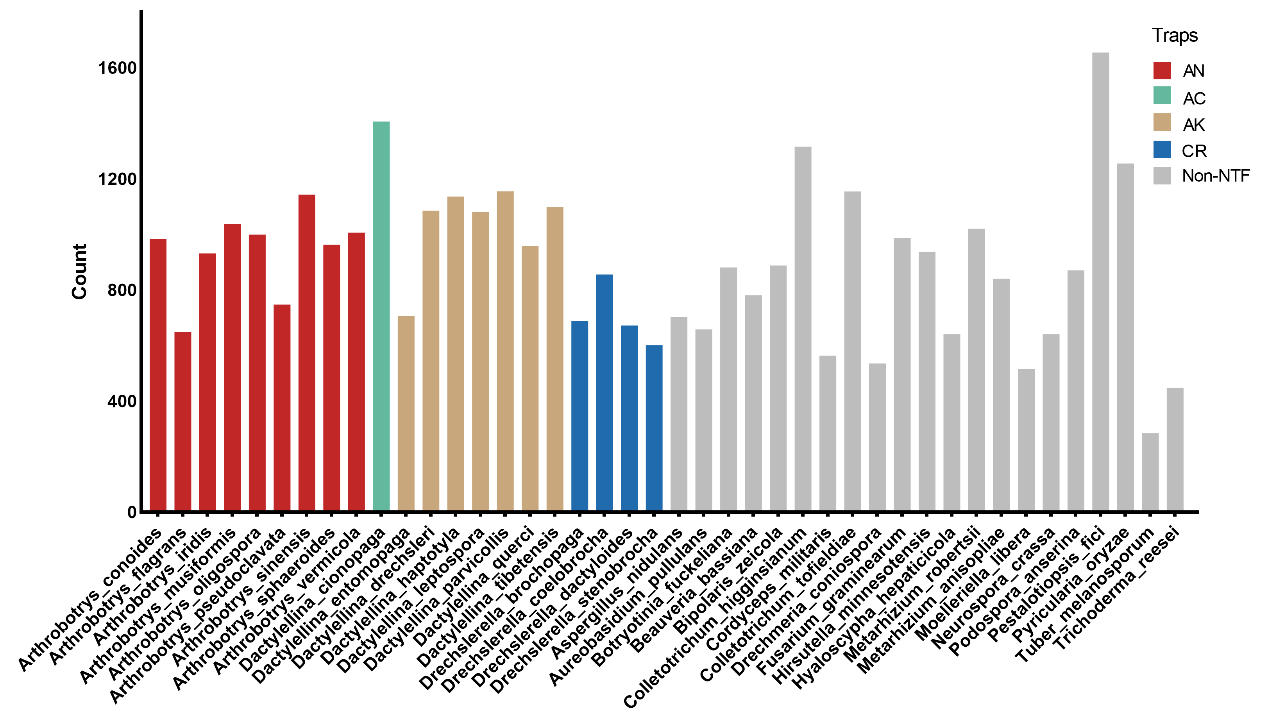

**Fig. S2:**  **The number of secreted protein genes in the 21 NTF and 21 non-NTF genomes.** Trapping devices formed by NTF are labelled: red (adhesive nets, AN), green (adhesive columns, AC), gold (adhesive knobs, AK), and blue (constricting rings, CR). Gray denotes non-NTF.

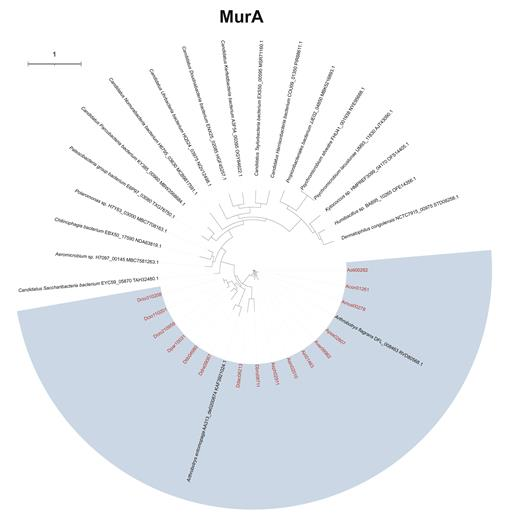

Fig. S3 cont.

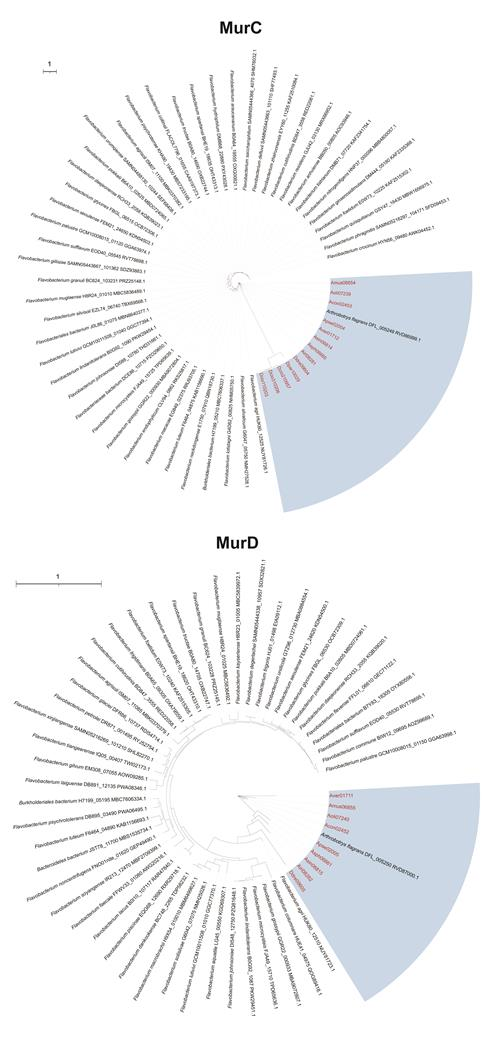

Fig. S3 cont.

**Fig. S3: Phylogenetic trees of three Mur proteins encoded by NTF and selected bacteria.** Maximum-likelihood trees based on the MurA, MurC and MurD protein sequences were constructed using RAxML. The bacterial species (only one genome sequence selected for each species) were chosen based on the top 100 BlastP hits when the corresponding *Arthrobotrys oligospora* protein sequences were used as queries. All NTF species (marked in red) are included in the blue background.

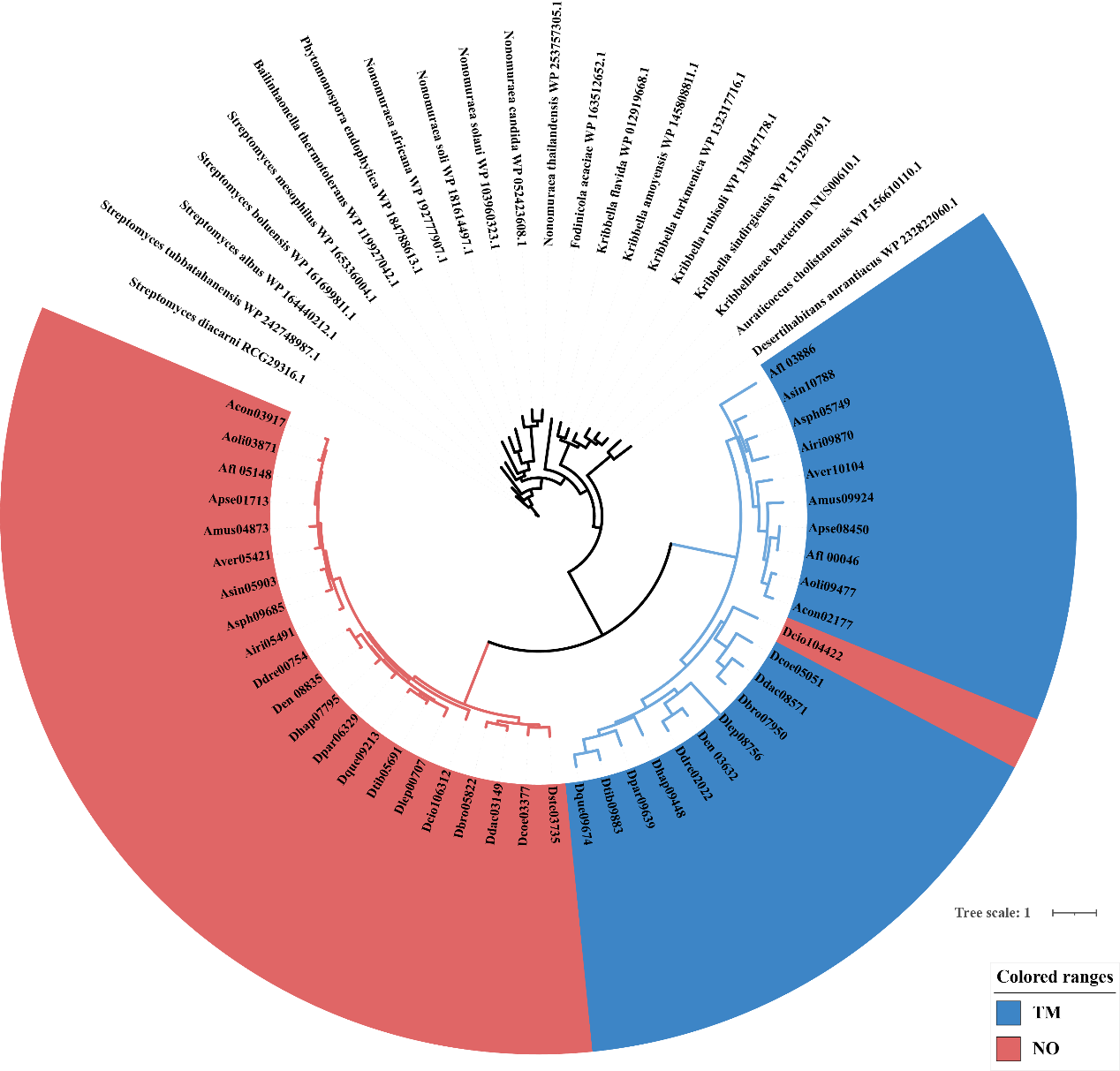

**Fig. S4: Maximum likelihood tree based on the Hyl proteins encoded by 21 NTF and selected bacteria.** Blue indicates those containing a transmembrane (TM) domain, and red signifies those lacking a TM domain.

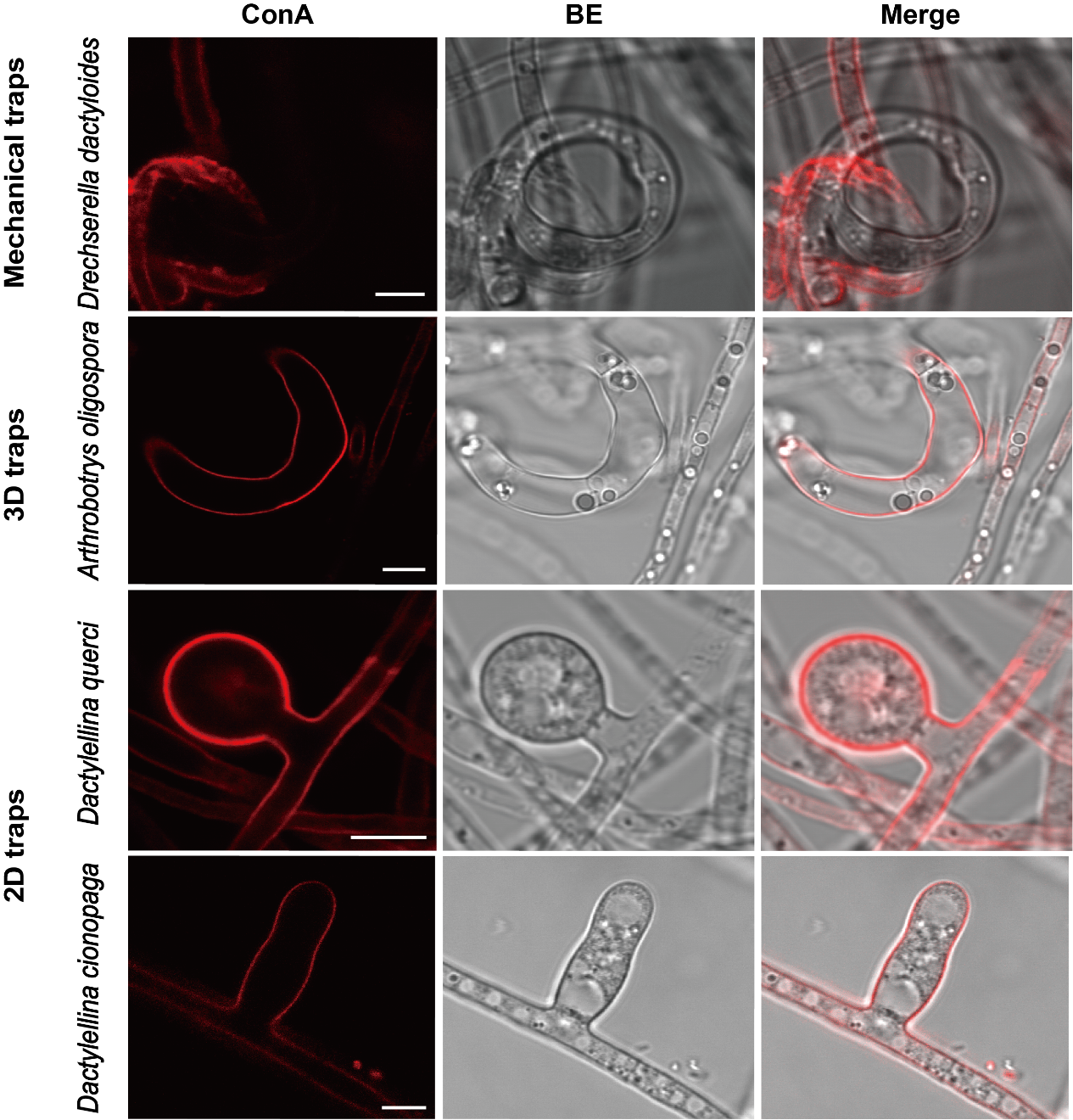

**Fig S5. Cell wall of NTF hyphae and traps stained by Concanavalin A (ConA).** ConA is a lectin that binds specifically to alpha-glucose and alpha-N-acetylglucosamine (GlcNAc) in the cell wall and displays red fluorescence when excited by 543 nm light. The cell wall of adhesive traps was stained more intensely than that of hyphae, whereas the opposite pattern was observed for those forming mechanical traps.
